# Injectable borax-loaded alginate hydrogels reduce muscle atrophy, modulate inflammation, and generate neuroprotection in the SOD1^G93A^ mouse model of ALS via activation of the IGF–Akt–mTOR axis pathway

**DOI:** 10.1101/2023.11.14.567052

**Authors:** Ana Rodriguez-Romano, Juan Gonzalez-Valdivieso, Laura Moreno-Martinez, Juan Francisco Vázquez Costa, Rosario Osta, Patricia Rico

**Affiliations:** Biomedical Research Networking Center in Bioengineering, Biomaterials and Nanomedicine (CIBER-BBN), Universitat Politècnica de València, 46022 Valencia, Spain; Center for Biomaterials and Tissue Engineering (CBIT), Universitat Politècnica de València, 46022 Valencia, Spain; Universidad de Valladolid, 47002 Valladolid, Spain; Biomedical Research Networking Center in Neurodegenerative Disorders (CIBERNED), University of Zaragoza, 50013 Zaragoza, Spain; Laboratory of Genetics and Biochemistry (LAGENBIO), Faculty of Veterinary-IIS Aragón, IA2-CITA, University of Zaragoza, Miguel Servet 177, 50013 Zaragoza, Spain; Biomedical Research Networking Center on Rare Diseases (CIBERER), Hospital Universitario y Politécnico la Fe, IIS La Fe, 46026 Valencia, Spain; Neuromuscular Unit, Hospital Universitario y Politécnico la Fe, IIS La Fe, 46026 Valencia, Spain

**Keywords:** Borax, ALS, muscle regeneration, NaBC1 transporter (SLC4A11), alginate hydrogel

## Abstract

Amyotrophic Lateral Sclerosis (ALS) is the most frequent and fatal condition that causes motor neuron loss and skeletal muscle paralysis. Although ALS is associated with mutations in over 40 genes, its etiology remains largely elusive without a cure or effective treatment. Historically considered the prototype of motor neuron diseases, ALS is defined today as a multisystem disorder that presents several changes in non-neuronal cell types, such as pathological changes in muscle occurring before disease onset and independent from motor neuron degeneration (dying back hypothesis). We base on the hypothesis that skeletal muscle may have an active contribution to disease pathology and thus we consider skeletal muscle tissue as a therapeutic target for ALS.

In previous works, we have demonstrated that boron transporter NaBC1 (encoded by the *SLC4A11* gene), after activation co-localizes with integrins and growth factor receptors producing a functional cluster that synergistically enhances crosstalk mechanisms accelerating muscle repair. In this work, we aimed to study the effects of borax (B) in a SOD1 mouse model of ALS targeting muscle. We have engineered and characterized injectable alginate-based hydrogels with controlled local borax release to effectively activate muscle NaBC1 *in vivo*. Treated mice presented improved motor function and extended survival correlated with the activation of essential muscle metabolic pathways, resulting in an enhanced muscle repair response and reduced muscle atrophy and inflammation. Interestingly, the activation of muscle repair mechanisms at the local level produced retrograde neuroprotection by motor neuron preservation and reduction in neuroinflammation. Altogether, this work presents evidence supporting the involvement of muscle tissue in ALS pathology, reinforcing skeletal muscle as a primary target to develop new therapies for ALS. We propose a novel strategy based on NaBC1 activation for ALS muscle regeneration.

**Graphical abstract:** 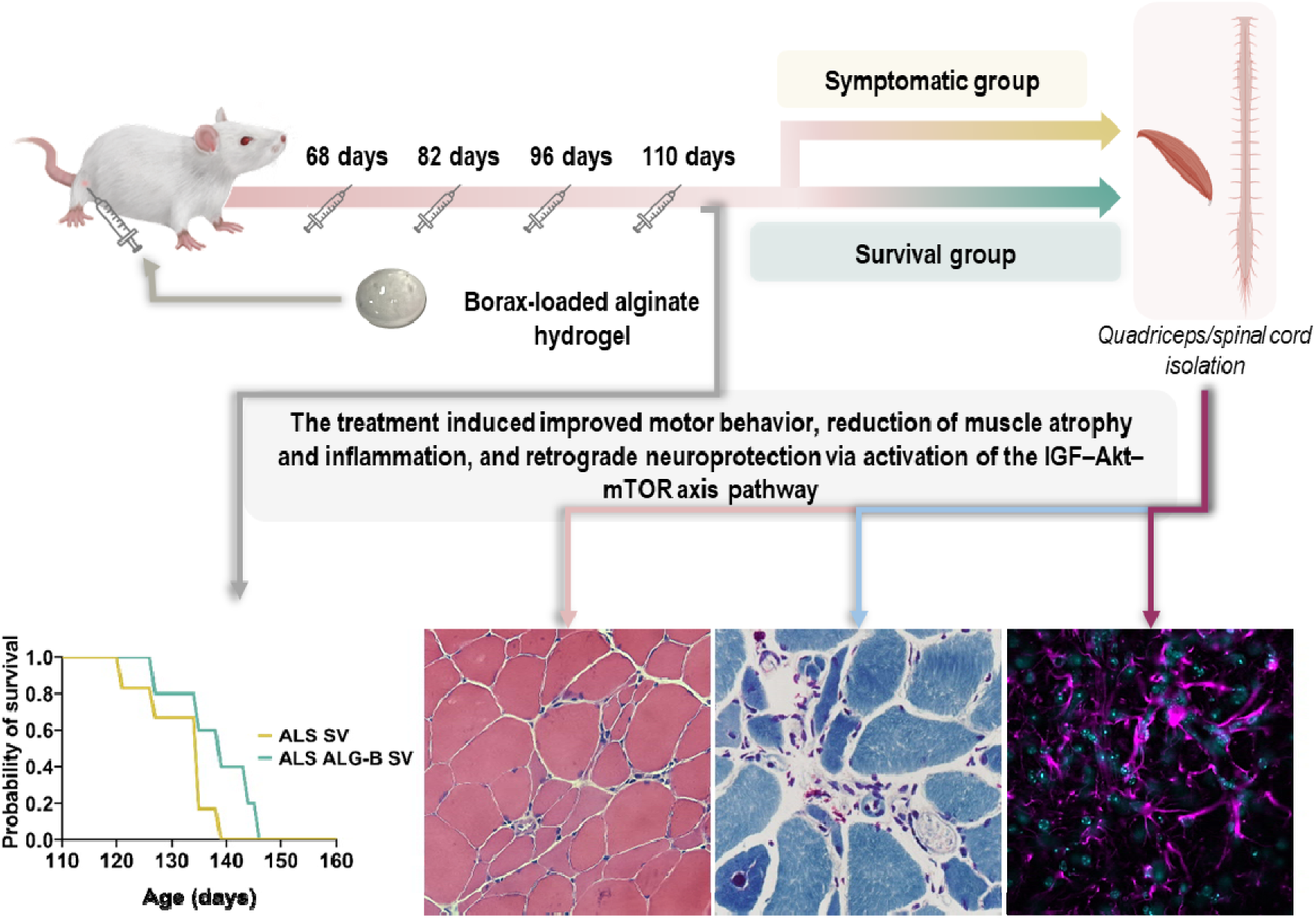

## 1. Introduction

Amyotrophic Lateral Sclerosis (ALS) is the most frequent and fatal entity affecting motor neurons.^[1]^ While the majority of ALS cases are sporadic, about 10% of the cases exhibit familial traits usually following an autosomal dominant inheritance. Clinical manifestations in sporadic and familial ALS are indistinguishable, suggesting the convergence of diverse pathways in neuromuscular degeneration. Despite the identification of over 40 genes associated with ALS (https://alsod.ac.uk/), the etiology of ALS remains elusive, with no cure or effective treatment available, only multidisciplinary care.^[2]^

The primary challenge in comprehending the pathogenesis of ALS lies in the remarkable clinical heterogeneity of the disease phenotype and course, even among patients harboring the same mutation.^[3,4]^ Additionally, the multisystemic nature of ALS pathology, spanning various biological systems, further complicates the identification of a suitable therapeutic target.

Although ALS is considered the prototype of motor neuron diseases, muscle weakness and degeneration is one of its clinic-pathological hallmarks since the disease was described by Charcot as “amyotrophic”. Recent evidence supports the view of ALS as a multisystem disorder involving changes also in non-neuronal cell types. A growing number of clinical and animal/cellular studies suggest that motor neuron damage may arise or be increased from mechanisms occurring in the glia^[5]^ or muscle cells.^[6]^ Moreover, the “dying back” hypothesis suggests that pathology originates in the skeletal muscle and is transmitted to the motor neuron before the onset of motor symptoms.^[7,8]^ Muscle-nerve interactions during muscle repair are essential to preserve muscle motor function, and skeletal muscle shows a key role in these events by secreting specific factors during regeneration. The observations of pathological changes in muscle preceding disease onset and independent of motor neuron degeneration highlight the significance of focusing on muscle tissue in ALS research and underscores its potential as a target for ALS therapies.^[9]^

Active therapies targeting motor neuron and muscle atrophy are based on the use of growth factors (GFs) either soluble,^[10]^ over-expressed in mouse models^[11]^, or stem cell-based growth factor delivery targeting glial cells^[12]^ and muscle inflammation.^[13]^ Nevertheless, only limited reports detail material systems designed for ion delivery as bioactive elements to activate intracellular signaling.^[14–17]^ Still there is a need to innovate and establish new methods and materials that facilitate the repair and functional regeneration of skeletal muscle.

Ions are capable of stimulating intracellular signaling, and some have potential as proangiogenic factors^[17]^ opening a new avenue for their application in tissue engineering. Muscle atrophy is also often related to defects in cytosolic calcium homeostasis due to oxidative stress, and thus targeting the calcium leakage from the muscle cells is another approach followed to improve muscle function.^[18,19]^ Boron plays a key role in mammalian cells in several metabolic pathways and in combination with other ions such as Ca^2+^ and Mg^2+^ as well as steroid hormones.^[20,21]^ Although boron is essential in the metabolism of mammalian cells,^[22]^ little is known about its homeostasis and function.

The borate transporter NaBC1 (encoded by the *SLC4A11* gene), controls boron homeostasis and functions as a Na^+^-coupled borate co-transporter.^[23]^ Recently, we have demonstrated for the first time a novel function for NaBC1. NaBC1 not only controls boron homeostasis, but also co-localizes with integrins and growth factor receptors (GFRs) after its activation, producing a functional cluster that synergistically enhances biochemical signals and crosstalk mechanisms enhancing vascularization,^[24]^ adhesion-driven osteogenesis,^[25]^ and muscle regeneration.^[26,27]^

In the present study, and based on our previous works, we investigated in a local proof of concept, the therapeutic efficacy of borax released from injectable alginate hydrogels targeting ALS muscle. We have used the SOD1^G93A^ mouse model which recapitulates lower motor neuron degeneration, and muscle denervation atrophy occurs before any evident sign of neurodegeneration.^[7,28]^ We found that only four injections of the treatment administered in the quadriceps of symptomatic mice, significantly improved motor function. This intervention also effectively reduced muscle atrophy, delaying evident symptom onset and extending mice survival. These positive outcomes were closely associated with muscle satellite cell activation and further progress through the myogenic program, crucial events for muscle regeneration and repair. Activation of NaBC1 potentiated muscle repair processes, resulting in the reduction of several pathological features characteristic of ALS-afflicted muscles and a decrease in muscle inflammation together with the activation of essential muscle metabolic pathways. Moreover, our data revealed that the recovery of muscle pathology had a retrograde neuroprotective effect, modulating neuro-inflammation and mitigating the loss of motor neurons. These findings play a crucial role in understanding the involvement of muscle tissue in ALS pathology, highlighting skeletal muscle as a primary therapeutic target. We propose a novel strategy based on the unique mode of action of active-NaBC1 to explore innovative treatments for ALS muscle regeneration.

## 2. Results and Discussion

### 2.1. Local borax-release preserves motor function, delays evident symptoms onset, and prolongs survival in a SOD1^G93A^ mouse model

In this work, we aimed to assess the effects of boron in the form of borax (B), as a local proof of concept, in an ALS murine model (B6SJL-Tg (SOD1^G93A^) 1 Gur/J ALS) to evaluate the possible effects of B in the recovery of the pathological conditions of muscle and nerve inflammation caused by ALS degeneration. To do that, we engineered and characterized injectable alginate-based hydrogels with controlled local B-release as previously described.^[26]^ We loaded the hydrogels by a simple mixing procedure that can be easily translated to the clinic, with borax 0.15 M equivalent to 6 mg per 100 µL hydrogel (ALG-B). **Figure S1-a** shows the non-cumulative B-release of 6 mm injectable hydrogels. Release from ALG-B hydrogels resulted in approximately 1,200 mg L^−1^ at day 1, despite the initial loading of 6,000 mg L^−1^ indicating that part of the borax content remains entrapped within the hydrogel and becomes sustained released over time. **Figure S1-b** illustrates the percentage of borax released corresponding to approximately 20 % of borax after 1 day.

Four subcutaneous injections in the quadriceps of 100 µL of either saline solution (ALS), ALG hydrogel (ALS ALG), or ALG B-loaded (ALS ALG-B), were administered to SOD1^G93A^ ALS mice, starting before visible symptoms onset (at 68 days old) and continuing every two weeks (at 82, 96 and 110 days old). Treated mice, male and female, were randomly distributed and organized in two parallel experiments: i) Symptomatic mice: three groups (*n* = 6), ALS control, ALS ALG, and ALS ALG-B, sacrificed at 115 days old for muscle and nerve evaluation independently of their clinical stage. ii) Symptomatic mice for survival (SV) assessment: two groups (*n* = 10), ALS SV control and ALS ALG-B SV, sacrificed at their final endpoint determined as the mousés inability to flip over within 30 seconds in the supine position determined as the final stage of disease progression-associated symptoms (ranging between 120-146 days).

For these experiments, changes in body weight and motor impairment were tracked using a behavioral four-limb hanging test over 12 weeks. One of the prominent characteristics of ALS in the SOD1^G93A^ mutant mouse model is its ability to induce severe muscle atrophy, leading to a progressive loss of weight due to a reduction in muscle mass and a significant decline in muscle strength.^[29]^ As depicted in **Figure 1-a**, the ALG-B group treated mice exhibited an increase in body weight over time and demonstrated the most favorable outcomes in the four-limb hanging test (Fig. 1-b) when compared to the non-treated control groups, suggesting preservation of motor function.

**Figure 1.**
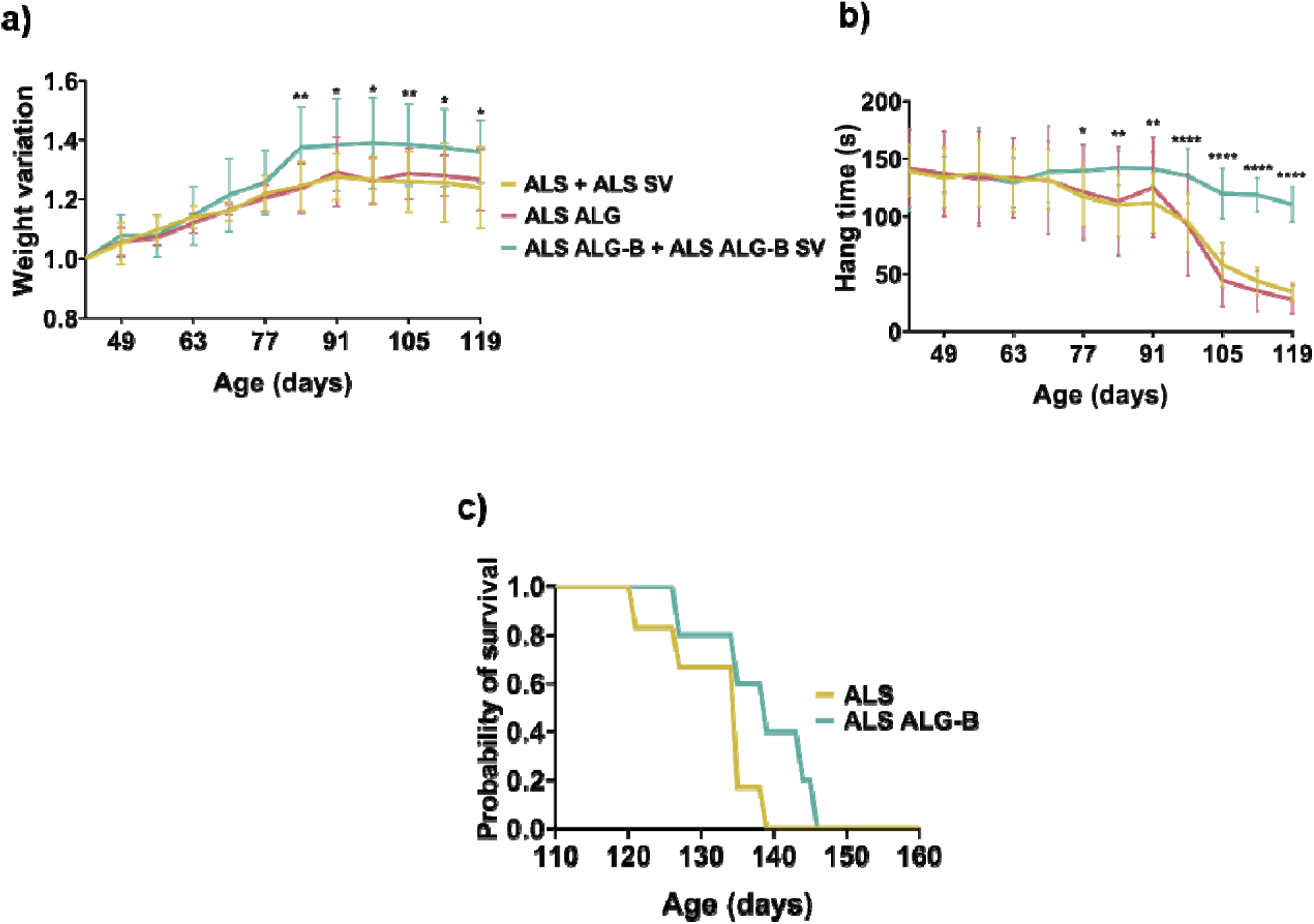
Evaluation of behavioral motor impairment and analysis of survival. a, b) Changes over time in weight and hanging time of mice (male and female) distributed in the different groups. Mice were subcutaneously injected with 100 µL of saline solution (ALS), ALG hydrogel (ALS ALG), or ALG B-loaded (ALS ALG-B) at 68, 82, 96 and 110 days old. Note that all the mice participating in the two parallel experiments, symptomatic mice, and terminal-stage mice for evaluation of survival (SV) were used for the measurements (ALS, *n* = 16; ALG, *n* = 4, ALG-B, *n* = 16). c) Kaplan-Meier survival curves of ALS (*n* = 10) and ALS ALG-B (*n* = 10) mice. Statistics are shown as mean ± standard deviation. For comparison between the three groups (ALS, ALG, ALG-B), data were analyzed by an ordinary one-way ANOVA test and corrected for multiple comparisons using Tukeýs correction analysis (p = 0.05). Statistics indicate differences between groups. ****p < 0.0001, **p < 0.01, *p < 0.05. For comparisons between the two groups (ALS SV, ALG-B SV), data were analyzed by an unpaired t-test applying Welch’s corrections (p = 0.05).

Mutant SOD1^G93A^ mice with elevated copy numbers generally maintain good health until around 80-90 days of age, even though at 60 days old they show altered muscle electromyography indicating muscle denervation. After 80-90 days old, they manifest visible symptoms of motor neuron disease, including limb tremors during suspension. By 120 days, most of these mice progress to advanced disease stages, resulting in complete paralysis.^[30]^ In addition to motor impairment, we monitored visible symptom progression in the survival (SV) experimental group. **Supplementary Table 1** illustrates the onset, endpoint, and duration of symptoms in each group. The group treated with ALG-B exhibited a non-statistically significant delayed onset of symptoms and an extended survival period (Fig. 1-c).

The observed improvement in motor function, along with the delay in visible symptoms onset and extended survival following local muscle treatment, suggests that muscle tissue may play a significant role in ALS pathology. This underscores the contribution of muscles to the pathological mechanisms affecting the disease, beyond motor neuron degeneration.

### 2.2. Local borax-release counteracts muscle atrophy in symptomatic and terminal-stage SOD1^G93A^ mice

After the euthanasia of symptomatic and terminal-stage mice, tissues were analyzed by histological techniques. The degeneration of muscle occurring after constant denervation and re-innervation processes, is characterized by progressive muscle atrophy, in which the fibers appear as angular atrophic fibers, surrounded by compensatory hypertrophic fibers to maintain homeostasis,^[31,32]^ and characterized by increased variability of fiber sizes, decreased fiber diameter and increased density of nuclear clumps between myofibers. For reliable quantification and measurement of pathology-relevant histological parameters of dystrophic muscle, we have followed TREAT-NMD standard operating procedures (SOP). **Figure 2** shows the histoarchitecture of muscle quadriceps analyzed in symptomatic (Fig. 2-a) and terminal-stage (Fig. 2-b) mice. **Figure S2** shows an illustration of the segmentation process performed for the quantification of complete muscle tissue sections.

**Figure 2.**
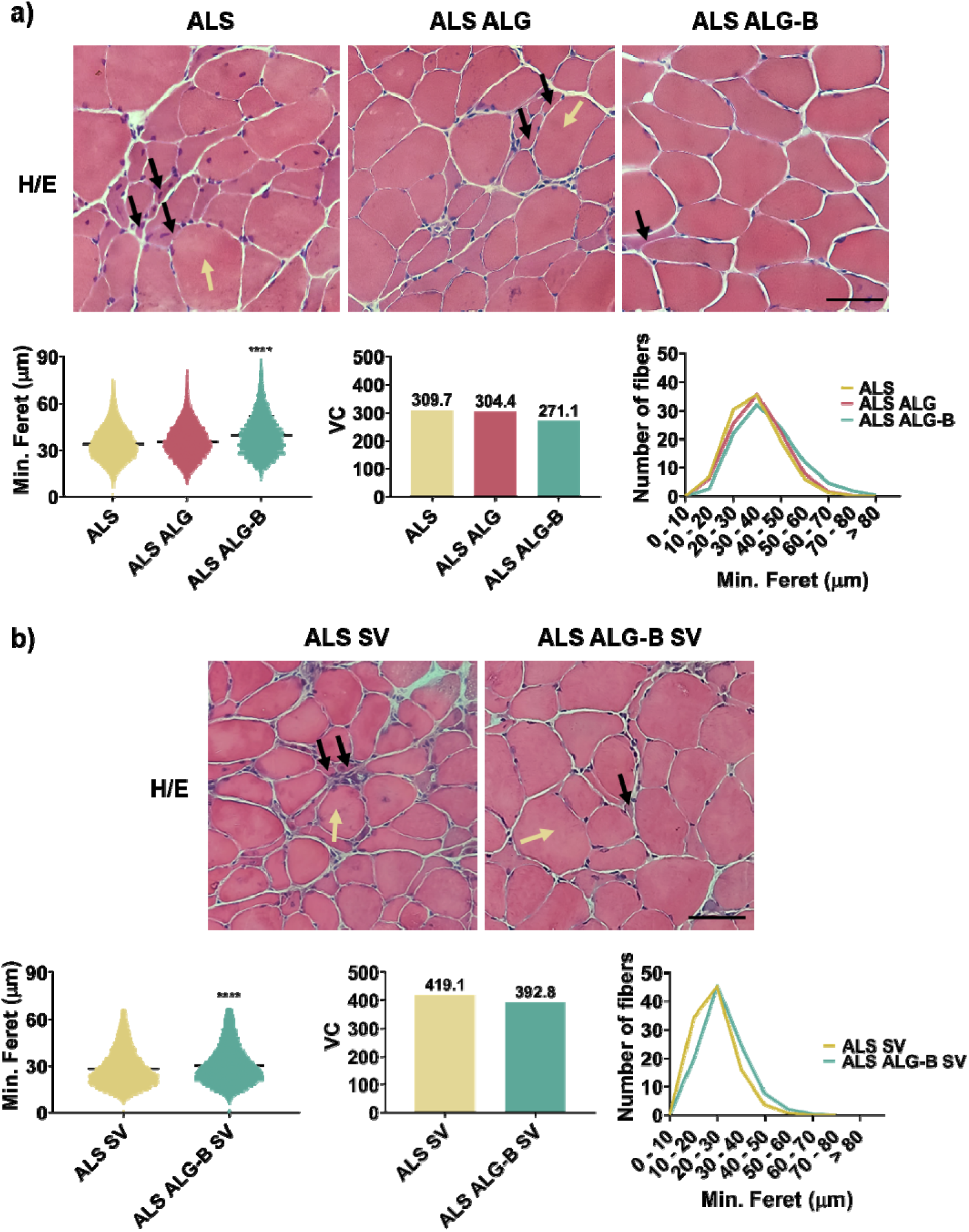
Morphometric muscle parameters in symptomatic and terminal-stage mice. Histoarchitecture of symptomatic (a) and terminal-stage (SV) (b) muscle quadriceps visualized by hematoxylin/eosin staining. In both animal experiments, the ALG-B group displayed reduced muscle atrophy, as evidenced by a decrease in atrophic fibers (10-20 µm, black arrows) surrounded by hypertrophic fibers (50-60 µm, yellow arrows). Scale bar 50 µm. Image analysis quantification of morphometric parameters including minimal Feret’s diameter and variability among fibers (Variance coefficient, VC). The ALG-B group exhibited a significant increase in the minimal Feret’s diameter and a decrease in VC, indicating a more uniform distribution of fiber sizes and the highest number of fibers with increased minimal Feret’s diameters. *n* > 25,000 fibers. Statistics are shown as mean ± standard deviation. For comparison between the three groups (ALS, ALG and ALG-B), data were analyzed by an ordinary two-way ANOVA test and corrected for multiple comparisons using Tukeýs correction analysis (p = 0.05). For comparisons between the two groups (ALS SV, ALG-B SV), data were analyzed by an unpaired t-test applying Welch’s corrections (p = 0.05). ****p < 0.0001.

For fiber size determination we have employed the Feret’s diameter of muscle fibers cross-sections. This is a robust morphometric parameter that avoids experimental errors related to fiber orientation and sectioning angle. Moreover, it reliably discriminates between dystrophic and normal phenotypes.

In both animal experiments, the ALS and ALG control groups displayed a decreased fiber size, a notable presence of atrophic fibers surrounded by hypertrophic fibers together with an increased density of nuclei, distinctly visible in the tissue sections (Fig. 2, arrows) and concordant with muscle atrophy, a pathological hallmark of ALS. We also observed the thickening of perimysium and endomysium, due to the augmented extracellular matrix components related to the appearance of muscle fibrosis. Despite muscle ALS pathologic changes being still evident, treated mice presented a significant increase in the minimal Feret’s diameter, suggesting an enlargement of muscle fibers. These results indicate the effective retardation of muscle atrophy progression by borax in both animal experiments and corroborate the treatment’s effectiveness in promoting muscle growth or preventing further muscle degeneration, as suggested by the improved muscle strength after motor behavior evaluation (Fig.1).

Furthermore, in both animal experiments, the ALG-B treated mice presented a reduction in the variability of fiber sizes, as expressed by the variance coefficient (VC), implying a decrease in the pathological characteristics present in ALS muscle. An analysis of the distribution of minimal Feret’s diameters revealed that the ALG-B group had the highest number of fibers with increased minimal Feret’s diameters. This finding is associated with an improved and accelerated regenerative response, indicating that the treatment is promoting muscle growth and function, and suggesting that some of the pathological features typically observed in ALS-afflicted muscles are mitigated.

The observation of reduced muscle atrophy in both symptomatic and terminal-stage mice suggests a lasting impact of borax treatment over time. Remarkably, just four injections of the treatment were sufficient to decelerate muscle atrophy progression, and these effects persisted until the end of the animals’ lives, even though the animals still exhibited ALS pathology and ultimately succumbed to the disease, likely due to the extensive neurological damage.

### 2.3. Local borax-release preserves fast (type II) muscle fibers, augments satellite Pax7^+^ cells, and reduces SOD1 and fibronectin levels in symptomatic and terminal-stage skeletal muscle of SOD1^G93A^ mice

Skeletal muscle is characterized by a heterogeneous composition of two types of fibers. Type I (slow-twitch) fibers, optimized for endurance activities, have a high content of myoglobin, essential for providing instant oxygen, and abundant mitochondria. Type II (fast-twitch) fibers are designed for short bursts of intense activity, have a major number of neuromuscular junctions, and contain large amounts of glycogen as a source for anaerobic glycolysis.^[33]^

In ALS, muscle denervation is accompanied by a progressive replacement of type II by type I fibers. This phenomenon, reported in both murine models and patients, highlights the vulnerability of fast-twitch fibers, which predominantly exhibit an atrophic appearance, while slow-twitch fibers adopt a compensatory hypertrophic morphology.^[31,32]^

Immunofluorescence analysis results of various muscle markers relevant to ALS is shown in **Figure 3**. Following borax treatment, we observed a substantial decrease in type I slow myofibers in the ALG-B muscle of both symptomatic and terminal-stage mice. Our findings suggest a potential preservation of type II fast myofibers. Given that type II fast fibers are the initial targets of atrophy in ALS, our results imply that the treatment is inhibiting the pathological myofiber transition characteristic of ALS. This aligns with our previous histological observations indicating a mitigation of muscle atrophy (Fig.2).

**Figure 3.**
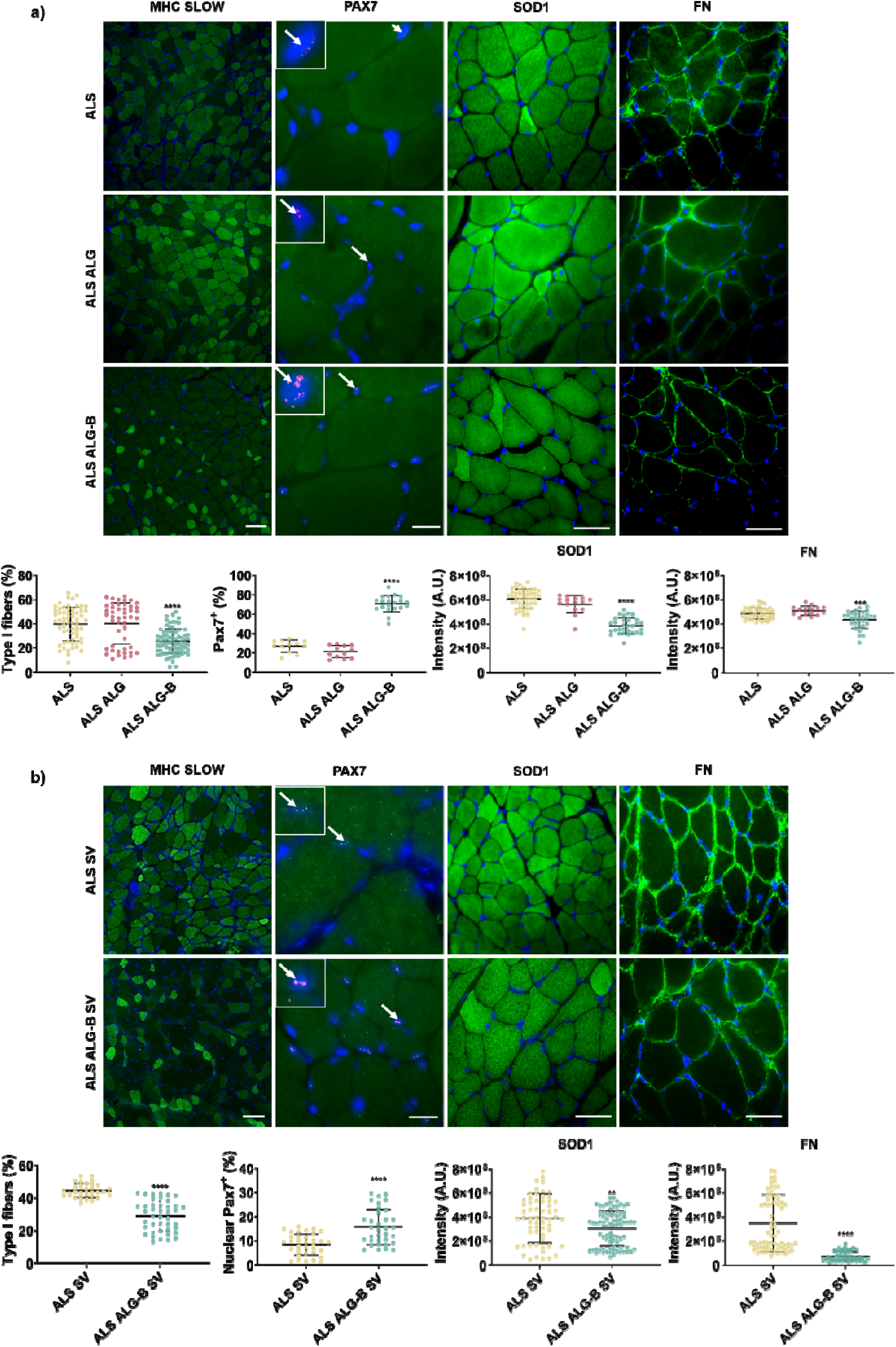
Immunofluorescence detection of ALS muscle markers in symptomatic and terminal-stage mice. Immunofluorescence detection of slow type I fibers, Pax7^+^ satellite cells, SOD1 protein aggregates, and fibronectin (FN). Quadriceps sections of symptomatic (a) and terminal-stage (b) mice were analyzed. In both animal experiments, the ALG-B group displayed reduced slow type I fibers, toxic SOD1 protein aggregates, and FN levels, in line with the observation of a reduction of muscle atrophy after the treatment and with the increase in Pax7^+^ cells. Scale bar 50 µm. Graphs represent the image analysis quantification of the intensity of the different markers. *n* > 12 images. Statistics are shown as mean ± standard deviation. For comparison between the three groups (ALS, ALG and ALG-B), data were analyzed by an ordinary two-way ANOVA test and corrected for multiple comparisons using Tukeýs correction analysis (p = 0.05). For comparisons between the two groups (ALS SV, ALG-B SV), data were analyzed by an unpaired t-test applying Welch’s corrections (p = 0.05). ****p < 0.0001, ***p < 0.001, **p < 0.01.

Several lines of evidence highlight the impact of ALS on muscle turnover, revealing that impaired myogenic processes could aggravate denervation-induced muscle wasting. Muscle denervation re-innervation cycles occurring in ALS, are accompanied by satellite cell activation (Pax7^+^ cells) that play an essential role in the intrinsic repairing mechanisms of muscle. ALS skeletal muscle satellite cells present impaired activity and myogenic potential. In ALS patients, Pax7 expression decreases^[32]^ and satellite cells become activated but do not progress through the myogenic program exhibiting incomplete differentiation and abnormal myotube and senescent-like morphology.^[34–36]^ In the SOD1^G93A^ ALS mouse model, satellite cells exhibit compromised cell proliferation capacity, even during the pre-symptomatic disease stages along with a reduction in the number of Pax7^+^ cells.^[37,38]^

Figure 3 shows a significant increase in Pax7^+^ cells in ALG-B treated mice as compared to control groups, evident in both symptomatic and terminal-stage mice. Notably, the percentage of Pax7^+^ cells in symptomatic ALS control mice is higher than in terminal-stage ALS control mice, underscoring the natural disease course and validating previously reported observations indicating a decline in Pax7^+^ cells throughout the disease.^[37,38]^ Given the reduced number and compromised proliferation of satellite cells in the SOD1^G93A^ model, our results indicating a substantial rise in active satellite cells suggest that borax is potentiating and enhancing muscle regeneration and repair processes. Together with the observed improvements in muscle strength and overall motor function (Fig.1), suggests that satellite cells are not only active, but they are also effectively progressing through the myogenic program reflecting an enhanced capacity for muscle healing.

Furthermore, the observed increase in Pax7^+^ cells found also in terminal-stage mice, who were in the advanced stages of the disease, indicates that treatment is offering long-term improvement potentially slowing down the progression of muscle degeneration.

We have also evaluated superoxide dismutase 1 (SOD1) and fibronectin (FN) expression at protein levels (Figure 3). SOD1 is a critical player in the anti-oxidative defense. The SOD1^G93A^ mouse model employed carries several copies of the human SOD1G93A mutation (glycine to alanine at position 93) which results in a ubiquitous toxic gain of SOD1 function.^[39,40]^ Specifically, skeletal muscle has been described as a primary target of SOD1^G93A^-mediated toxicity on which oxidative stress triggers muscle atrophy.^[29]^

ALG-B-treated symptomatic and terminal-stage mice presented a reduction in total SOD1 protein levels indicating that treatment is helping to protect muscle from toxic SOD1 oxidative stress effects slowing down muscle deterioration, and conferring an improved muscle function and strength, which translates into enhanced motor function observed in Fig.1 and reduction in muscle atrophy (Fig. 2). Further investigation is needed to decipher the specific mechanism by which borax reduces SOD1 protein levels, even though our results suggest that ALG-B treatment is specifically targeting SOD1 within the muscle and could represent a promising therapeutic approach.

In addition, after muscle denervation processes, augmented FN protein levels have been reported in both ALS patients^[41]^ and SOD1^G93A^ mice,^[42]^ related to the appearance of fibrotic changes. An active fibrotic process has been reported in the skeletal muscle of symptomatic SOD1^G93A^ mice.^[42]^

FN levels were also diminished in symptomatic and terminal-stage mice after treatment concordant with the observation of a reduced thickness of perimysium and endomysium (Fig.3), suggesting a reduction in muscle fibrosis levels related to FN overexpression.

Overall, our findings at the protein level in muscle histopathology, align with the observed recoveries in motor function (Fig. 1) and the deceleration of muscle atrophy progression (Fig. 2) indicating that local borax treatment is exerting a positive impact on muscle integrity and function.

### 2.4. Evaluation of muscle inflammation indicates that muscle borax-release reduces mast cell inflammatory response in symptomatic and terminal-stage SOD1^G93A^ mice

In ALS, inflammation is a key mechanism contributing to the degeneration process of both motor neurons and muscles. Several reports describe the presence of inflammatory cells within peripheral nerves and skeletal muscles of ALS patients and rodents.^[43,44]^ Among inflammatory cells, the granulated hematopoietic-derived mast cells, activate upon tissue damage and orchestrate the inflammatory responses recruiting and activating other immune cells through degranulation and release of inflammatory mediators and enzymes,^[45–48]^ mediating neurogenic inflammation.^[49]^ Recent findings suggest that mast cells directly interact with degenerating motor nerve endings and motor endplates in the skeletal muscle of SOD1^G93A^ rodents with their number and degranulation pattern correlating with the progression of paralysis,^[44]^ highlighting their involvement in the chronic inflammatory muscle response in ALS.^[50,51]^

The assessment of muscle mast cell recruitment revealed that in symptomatic mice (**Figure 4-b**), there was an elevation in the total number of infiltrated mast cells in the control groups (ALS, ALG), along with an increase in degranulated mast cell count and mast cell proximity to nerves. However, the ALG-B treated group showed a significant reduction in the density of infiltrated mast cells as well as the de-granulating mast cell number, suggesting an inhibition of mast cell migration and activation. Further, in B-treated quadriceps mast cells were mainly perivascular located and not near nerve fibers, suggesting that borax may downregulate mast cell-mediated inflammatory events influencing the degeneration of the muscle and the peripheral motor pathway.

**Figure 4.**
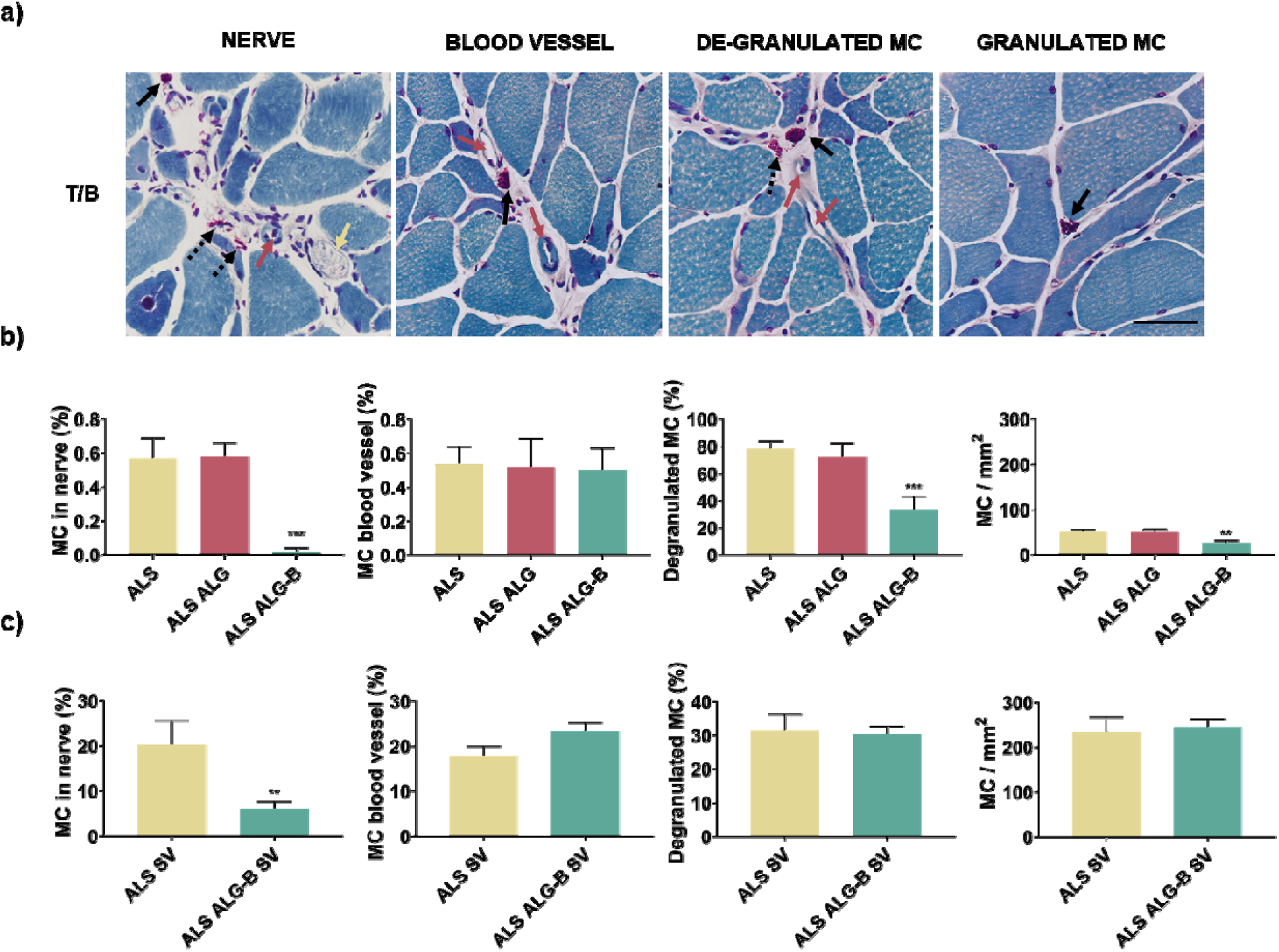
Infiltration and degranulation of mast cells into the skeletal muscle of symptomatic and terminal-stage SOD1^G93A^ mice. (a) Analysis of mast cell recruitment in muscle by toluidine-blue staining. Quadriceps sections of symptomatic (b) and terminal-stage (c) mice were analyzed. In symptomatic mice, the ALG-B group presented a strong decrease in the density of infiltrated mast cells as well as the de-granulating mast cell number. Further, mast cells were mainly perivascular located and not close to a nerve fiber, a hallmark characteristic of extensive nerve inflammation occurring in SOD1^G93A^ mice. In terminal-stage mice, the ALG-B group presented a strong decrease in mast cells in the nerve periphery, suggesting a delay in the inflammation onset despite the active inflammation process. Scale bar: 50 µm. Graphs represent the image analysis quantification of the percentage of mast cells in each location, the total number of mast cells, and the total number of de-granulated mast cells. *n* > 40 images. Statistics are shown as mean ± standard deviation. For comparison between the 3 groups (ALS, ALG, ALG-B), data were analyzed by an ordinary one-way ANOVA test and corrected for multiple comparisons using Tukeýs correction analysis (p = 0.05). For comparisons between 2 groups (ALS SV, ALG-B SV), data were analyzed by an unpaired t-test applying Welch’s corrections (p = 0.05). ***p < 0.001. **p < 0.01.

In terminal-stage mice (Fig. 4-c), we found high levels of total and degranulated mast cells, even in treated ALS ALG-B mice, indicating an ongoing pro-inflammatory process consistent with the disease’s advanced progression. Consequently, in terminal-stage mice, the total number of mast cells surrounding nerve fibers in control groups were higher compared to symptomatic control mice, reflecting the extensive nerve inflammation characteristic of the SOD1 mouse model (Figure 4-b,c). Nevertheless, despite the pronounced inflammation present in the terminal-stage ALS muscle, ALG-B treated mice exhibited a significant reduction in mast cells in the nerve periphery. This suggests that while inflammation does occur at a later stage, the treatment appears to delay its onset.

We have further analyzed IL-6 muscle levels, a cytokine known for its diverse functions and crucial role in immune response, inflammation, metabolism, and hematopoiesis. When the production of IL-6 becomes dysregulated, it can lead to the development of various chronic inflammatory diseases.^[52]^ This cytokine is produced by immune and blood cells, endothelial cells, and myocytes during muscle contraction.^[53]^ Interestingly, elevated levels of IL-6 have been observed in ALS patients, which is associated with motor neuron degeneration.^[54]^ Additionally, IL-6 has been found to contribute to muscle atrophy after denervation, further highlighting the involvement of cytokines in the progression of ALS. Given its significance, IL-6 has emerged as a potential biomarker for ALS and a target for therapeutic interventions, as demonstrated by ongoing clinical trials.^[55]^

**Figure S3** illustrates the immunofluorescence detection and image analysis quantification of IL-6 in symptomatic and terminal-stage mice. In both experiments, the control mice exhibited elevated levels of IL-6. Notably, the IL-6 levels were significantly higher in terminal-stage control mice, consistent with previous findings that associate IL-6 with disease progression.^[55]^ However, the quadriceps muscles treated with borax (ALG-B) presented reduced IL-6 levels, indicating a decrease in the inflammatory response within the muscle tissue following treatment. These findings collectively suggest that borax may function as a therapeutic agent by promoting muscle repair processes, suppressing mast cell-mediated inflammatory events, reducing pro-inflammatory cytokines such as IL-6, and mitigating muscle degeneration and neuroinflammation.

### 2.5. Local muscle borax-release induces motor neuron preservation and reduces glial inflammation in the spinal cord in symptomatic and terminal-stage SOD1^G93A^ mice

Following the assessment of muscle inflammation, which revealed diminished inflammation specifically near muscle nerve fibers (Fig.4, S3), and the observed changes in improved motor behavior (Fig.1) and improved muscle health (Fig.2, 3) that indirectly can potentially be associated with enhanced motor function or survival, we aimed to determine if the administration of a localized muscle treatment to ALS SOD1^G93A^ mice could lead to neuroprotection. To do that we analyzed the complete spinal cord of symptomatic and terminal-stage SOD1^G93A^ mice.

The SOD1^G93A^ model is characterized by motor neuron degeneration caused by the toxic effects of the SOD1 mutant protein, which affects both motor neurons and their neighboring glia.^[39,56]^ In **Figure 5**, our findings revealed a notable and significant increase in the number of motor neurons in the spinal cord of both symptomatic and terminal-stage mice following local muscle treatment compared to the control groups. These results suggest that the treatment applied to the muscle tissue may have broader effects on the central nervous system in ALS, potentially influencing the neurodegenerative process and preserving motor neurons beyond the targeted muscle area. Our results are in line with previous studies indicating that preservation of muscle tissue function can slow motor neuron degeneration,^[57]^ despite overexpression of mutant SOD1 in rodents induces inevitably a neuropathic phenotype. Furthermore, we conducted measurements to quantify the white matter area in the complete spinal cord. Our results indicated that both, symptomatic and terminal-stage mice treated with ALG-B exhibited also an increased white matter area, indicating the preservation of neuronal axon integrity. These findings align with the observed reduction in inflammation shown in Figures 4 and S3.

**Figure 5.**
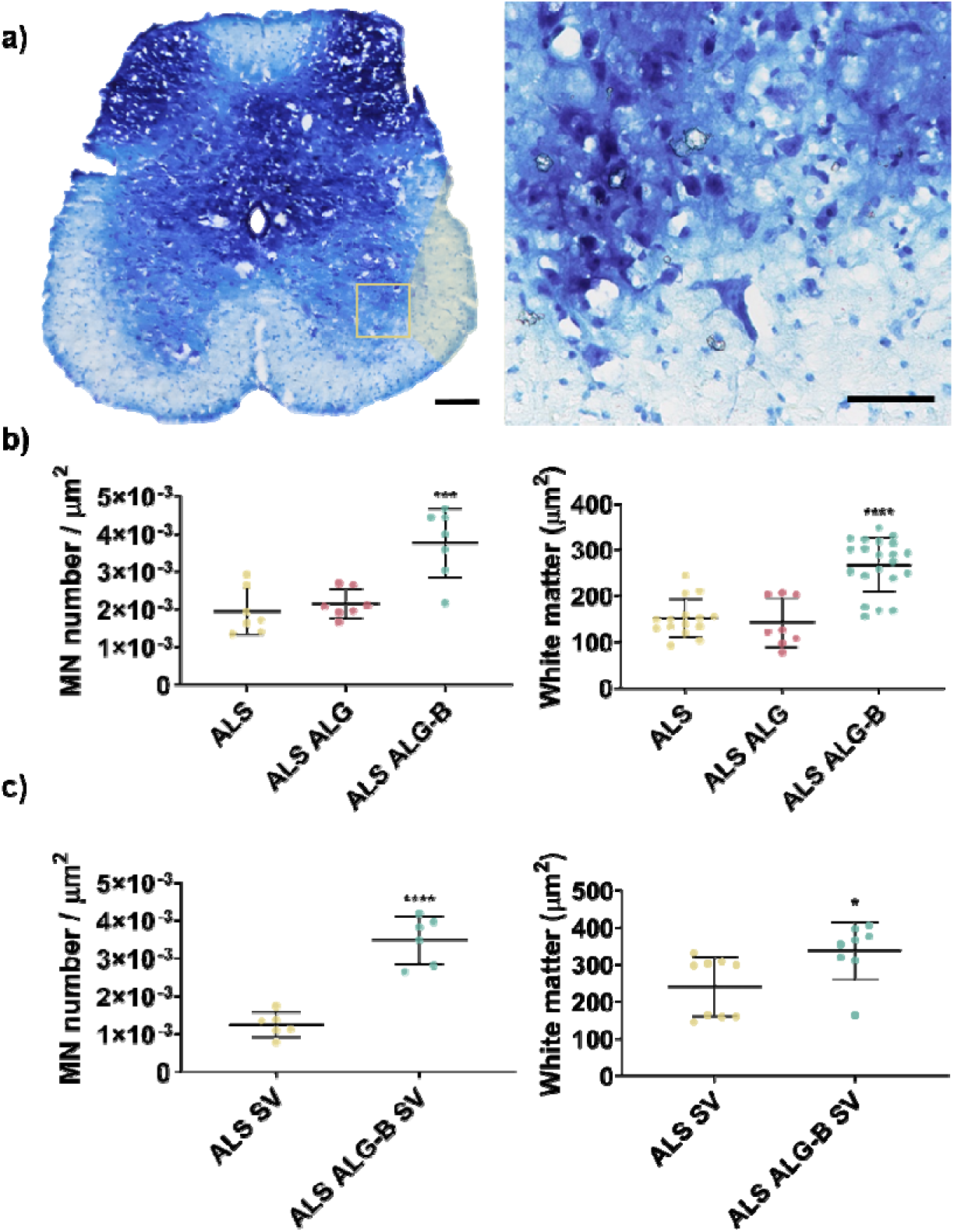
Evaluation of motor neuron preservation in spinal cord in symptomatic and terminal-stage SOD1^G93A^ mice. a) Representative image of spinal cord stained with toluidine-blue (scale bar: 250 µm) and detail of motor neuron located in the ventral horn (scale bar: 50 µm). Motor neuron count and white matter area in the spinal cord were assessed in symptomatic (b) and terminal-stage (c) mice. The spinal cord was segmented into caudal, dorsal, and cervical regions, and multiple histological sections from each segment were analyzed. The B treatment applied in muscle induced an increase in motor neuron number and white matter area in the spinal cord in both groups. *n* ≥ 6 mice per condition. Statistics are shown as mean ± standard deviation. For comparison between the three groups (ALS, ALG, ALG-B), data were analyzed by an ordinary one-way ANOVA test and corrected for multiple comparisons using Tukeýs correction analysis (p = 0.05). For comparisons between 2 groups (ALS SV, ALG-B SV), data were analyzed by an unpaired t-test applying Welch’s corrections (p = 0.05). ****p < 0.0001, ***p < 0.001, *p < 0.05.

In addition to the selective degeneration of motor neurons, astrogliosis is a prominent characteristic of ALS. Astrocytes, a type of glial cell, maintain neuronal homeostasis and provide support and protection for efficient neuronal function. However, in pathological conditions, activated astrocytes can have harmful effects on neuronal survival. Glial fibrillary acidic protein (GFAP) is an intermediate filament protein that is highly expressed in reactive astrocytes and is significantly upregulated in the spinal cord of ALS patients and mouse models, where reactive astrogliosis is prominent.^[58]^ The increased production of GFAP has been linked to modulatory roles in the progression of ALS.^[59]^ In **Figure 6**, we examined the detection of GFAP in the complete spinal cord of both symptomatic and terminal-stage SOD1^G93A^ mice. Similar to the observations made regarding mast cells and IL-6 levels in muscle tissue (Figures 4, S3), we found that neuro-inflammatory levels were significantly higher in terminal-stage mice, corroborating that the GFAP protein gradually increases with disease progression, consistent with previous reports.^[57]^ Interestingly, following the local muscle treatment, we observed a reduction in GFAP values in the treated mice from both experiments. These findings indicate that the application of borax in muscle has the potential to modulate neuroinflammation processes that take place in ALS. However, it is important to note the possibility of a direct impact of borax on glial cells and motor neurons. Additional research is necessary to unravel the precise cellular-level function of borax and NaBC1 in the nervous system.

**Figure 6.**
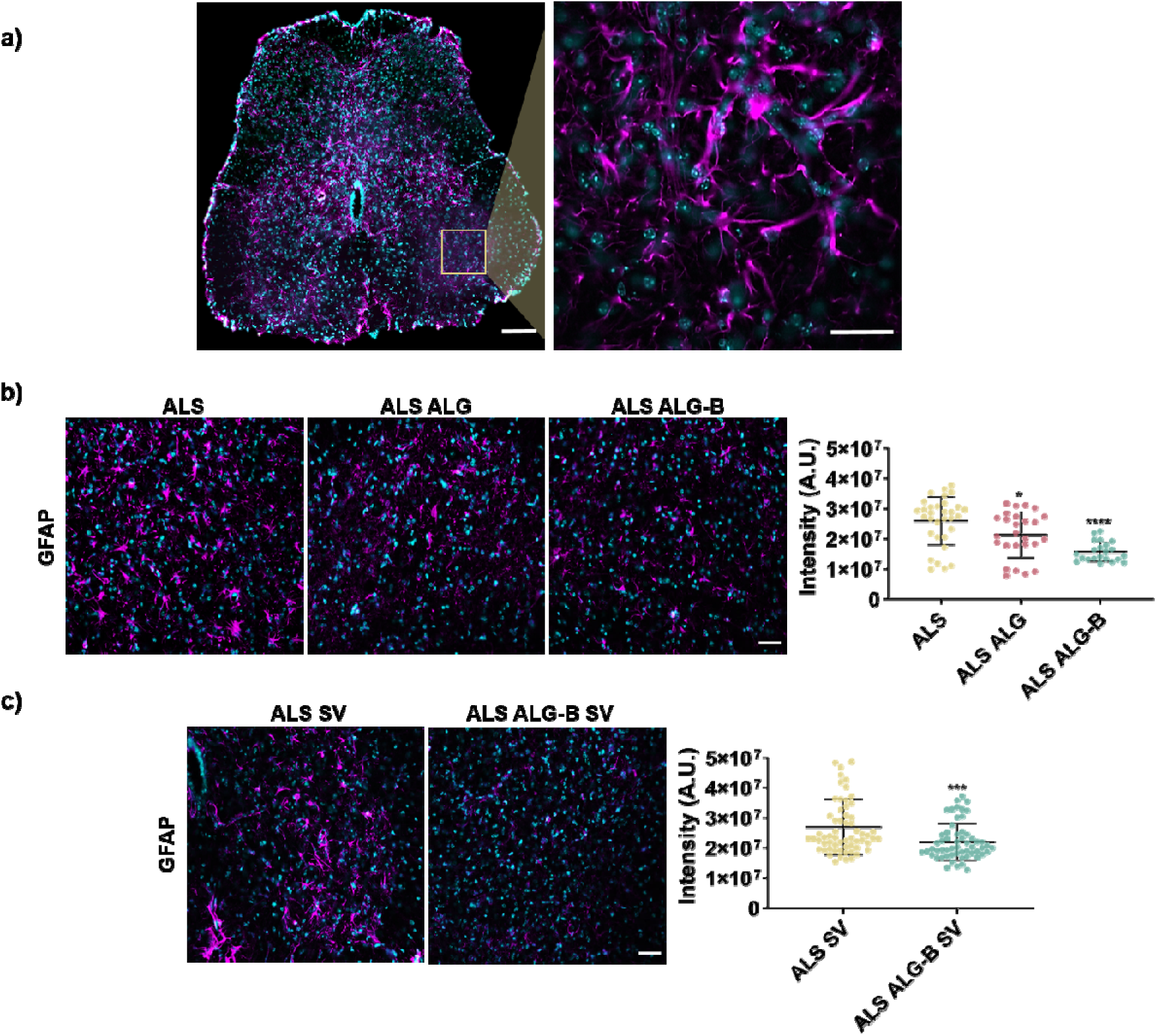
Evaluation of glial-mediated inflammation in the spinal cord in symptomatic and terminal-stage SOD1^G93A^ mice. a) Representative image of the spinal cord caudal region (scale bar: 250 µm), with detailed ventral horn area where astrocytes are detected with GFAP antibody (magenta, scale bar: 25 µm). Cellular nuclei were detected with DAPI. Immunofluorescent detection of GFAP in symptomatic (b) and terminal-stage (c) mice. Scale bar: 50 µm. ALG-B treatment decreases GFAP expression, indicating a reduction of neuroinflammation. *n* **>** 30 images per condition. Statistics are shown as mean ± standard deviation. For comparison between the three groups (ALS, ALG, ALG-B), data were analyzed by an ordinary one-way ANOVA test and corrected for multiple comparisons using Tukeýs correction analysis (p = 0.05). For comparisons between the two groups (ALS SV, ALG-B SV), data were analyzed by an unpaired t-test applying Welch’s corrections (p = 0.05). ****p < 0.0001, ***p < 0.001, *p < 0.05.

### 2.6. Analysis of muscle gene expression in symptomatic and terminal-stage SOD1^G93A^ mice

After the euthanasia of symptomatic and terminal-stage mice, quadriceps muscles were used for RNA extraction and analysis of gene expression (**Figure 7**).

**Figure 7.**
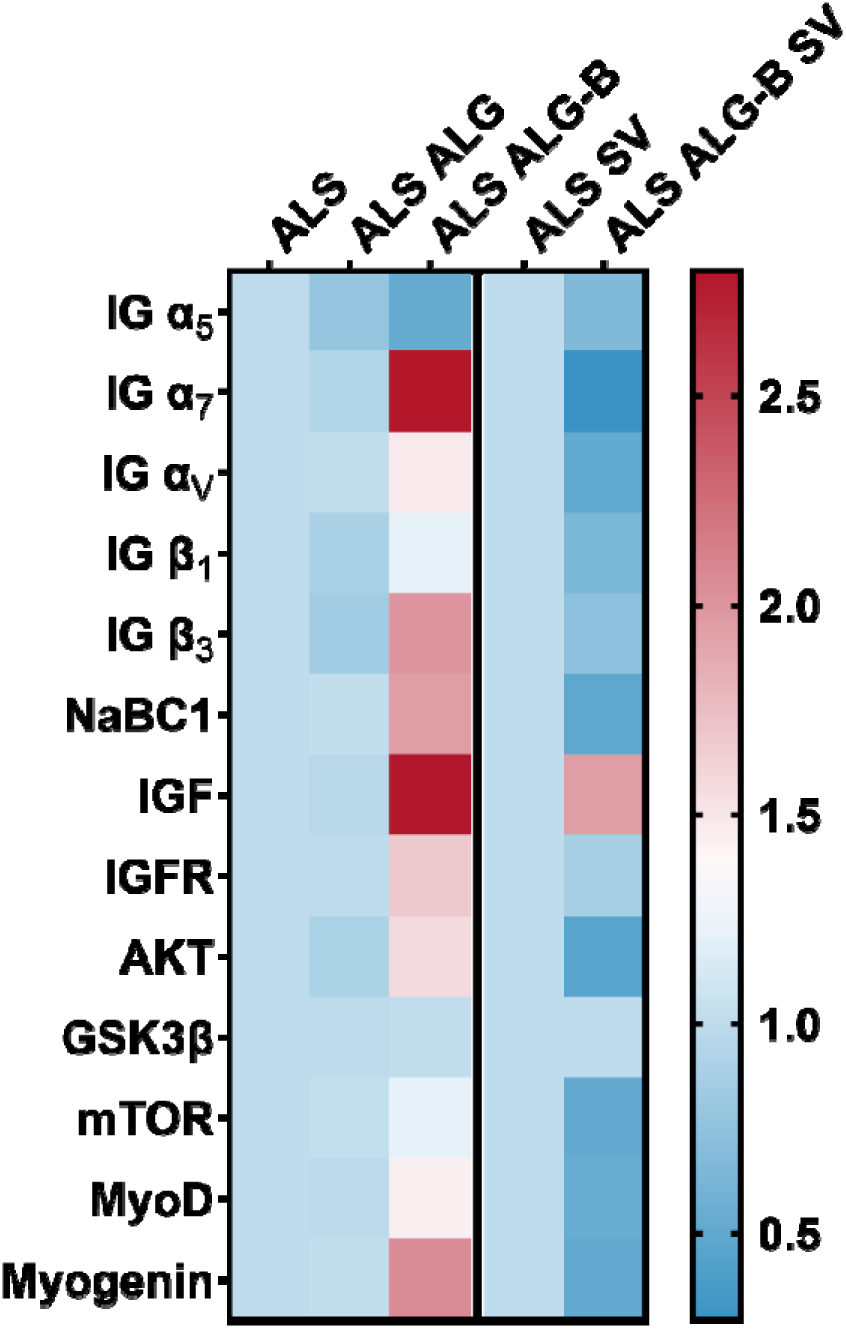
Representative heat map analysis of the up/down-regulated genes in the muscle of symptomatic and terminal-stage SOD1^G93A^ mice. Quantitative real-time PCR analysis of relative mRNA expression of α_5_, α_7_, α_v_, β_1_ and β_3_ integrins, NaBC1 transporter, members of the IGF-1–Akt–mTOR axis involved in muscle growth, and myogenic regulatory factors (MyoD, Myogenin). *n* ≥ 6 mice per condition.

In our previous works, we have shown *in vitro* that NaBC1 co-localizes with FN-binding integrins (α_5_β_1_ and α_v_β_3_) after activation, and cooperates with GFR stimulating intracellular signaling.^[24–26]^

The skeletal muscle extracellular matrix (ECM) is a dynamic structure primarily comprised of laminins (LM), collagens, proteoglycans, and FN. This intricate network plays a pivotal role in regulating the physiological functions of muscles.^[60,61]^ Specifically, FN holds a crucial position in muscle regeneration by aiding in the proliferation and differentiation of satellite cells.^[62]^ Moreover, FN-binding integrins such as α_5_β_1_ and α_v_β_3_ contribute significantly to muscle stability and exert a regulatory influence on the repair processes.^[63,64]^ Within the skeletal muscle tissue, integrins form connections with the ECM, facilitating interactions among muscle cells within their environment. These interactions are vital for muscle cell adhesion, migration, and differentiation.^[65]^

In this work, aiming to confirm the interplay between NaBC1 and integrins for muscle recovery *in vivo*, we have evaluated FN-binding (α_5_β_1_ and α_v_β_3_) and LM-binding (α_7_β_1_) integrins expression. α_7_β_1_ integrin, is specifically associated with skeletal muscle playing a key role in muscle stability, function, and repair^[66]^ and acts as a critical mechanosensor contributing to mechanical-load induced skeletal muscle growth.^[67]^

Our results showed that symptomatic mice treated with ALG-B hydrogels presented a markedly upregulated mRNA expression of α_7_, α_v_, β_1_ and β_3_ integrins, ranging from a 1.2 to 2.8-fold rise (Fig.7), in comparison to the controls. This indicates a stimulation of muscle repair and regeneration after muscle local treatment. Particularly noteworthy was the substantial 2.8-fold increase in mRNA expression of α_7_integrin. Given the pivotal role played by α_7_β_1_ integrin in skeletal muscle ECM attachment and muscle force development^[68]^, the absence of this integrin causes muscle defects and myopathy.^[69]^ Moreover, elevating α_7_ levels has been employed as a strategy to counterbalance deficiencies in ECM and cytoskeleton linkage occurring in certain muscular dystrophies, leading to a reduction in muscle pathology.^[70]^ Thus, our results showing induced upregulation of integrin expression, and specifically α_7_, after NaBC1 activation, reinforce our hypothesis that considers NaBC1 and integrin stimulation for enhancing the intracellular signaling as we previously reported.^[24–26]^

Interestingly, despite our previous observations of upregulated α_5_ integrin expression *in vitro* during cell adhesion stages following NaBC1 activation in healthy murine muscle cells,^[26]^ our current findings show decreased levels of this integrin *in vivo* in symptomatic SOD1^G93A^ mice. Specifically in ALS, α_5_ integrin has been shown to be upregulated in the spinal cord and peripheral nerves, contributing to immune cell recruitment and inflammation, factors linked to disease progression.^[71]^ In fact, α_5_ has been proposed as a potential therapeutic target since α_5_ blocking resulted in improved motor function, delayed disease progression, and increased survival in ALS animal models.^[71]^ Figure 7 shows that treatment with borax-loaded hydrogels halved the expression levels of α_5_ integrin in symptomatic mice. Taken together the obtained results describing a reduction of inflammation in spinal cord (Fig. 6) and muscle (Figures 4, S3), we underscore the potential of borax as a therapeutic approach for the treatment of ALS.

We have also observed that mRNA expression of NaBC1 transporter was upregulated in ALG-B symptomatic mice, suggesting a concentration-dependent regulatory mechanism exerted by borax in myotubes for NaBC1 expression. To date, no data are available regarding NaBC1 expression regulation in mammalian cells.

Additionally, we have assessed key metabolic pathways that impact muscle homeostasis and function. The Insulin-like growth factor (IGF) stimulates myogenesis, proteostasis and cell growth through the Akt/mTOR signaling pathway.^[72]^ This pathway controls various cellular functions, including cell growth and proliferation.^[72]^ Furthermore, mTOR plays a role in regulating metabolism and energy equilibrium by managing nutrient detection and utilization within muscle tissue.^[73]^ In our study, we have observed increased mRNA expression of IGF and its receptor, IGF receptor (IGFR), leading to increased Akt and mTOR levels in symptomatic mice treated with ALG-B. Moreover, our findings indicated a subsequent decrease in GSK3β in these mice, aligned with the inhibition of GSK3β due to Akt activation, as these pathways oppose each other metabolically. Intriguingly, GSK3β has been observed to be overexpressed in non-neural cells of ALS patients, suggesting its potential involvement in the disease’s progression.^[74]^ Further, GSK3β hyperactivity has been confirmed as a causative factor in disease pathogenesis, while its inhibition is a potential therapeutic avenue.^[75,76]^ Our results demonstrate that active NaBC1 triggers intracellular signaling via activation of IGF-Akt-mTOR axis. From a therapeutic perspective, considering that IGF has been used as a muscle neurotrophic factor to counteract impairment of the proteostasis observed in ALS^[77]^ we propose that NaBC1 activation represents a promising new therapeutic avenue to address impaired muscle physiology in ALS.

We have further evaluated two members of the Myogenic Regulatory Factor (MRF) family of transcription factors, MyoD and Myogenin. MyoD is a master gene for myogenesis. Activated satellite cells expressed MyoD at early stages of myogenic differentiation and is crucial for efficient activation and proliferation.^[77,78]^ After MyoD activation in turn induces expression of other myogenic-related genes such as Myogenin. Myogenin participates in myocyte fusion and is essential for regulation of adult myofiber growth and satellite cell homeostasis.^[79]^ Our results further corroborated the borax restorative effects in muscles of symptomatic mice showing upregulation of MyoD and Myogenin (1.5 and 2.1-fold increase respectively), and are in line with the observations confirming improvement of motor function (Fig.1) and muscle atrophy recovery (Fig.2). However, the results obtained in terminal-stage mice presented a contrasting outcome, displaying a general downregulation of all evaluated genes. We attribute this outcome to the severe damage evident in the muscles of these mice, stemming from the extensive neurodegeneration and muscle denervation process occurring in ALS during the disease’s advanced stages. These terminal-stage mice were sacrificed after reaching total paralysis, rendering them unable to self-feed or drink. Despite displaying noticeable improvements in muscle histoarchitecture, particularly in muscle atrophy recovery and inflammation reduction, these mice ultimately exhibited minimal muscle metabolic activity, likely due to energy depletion from starvation and the absence of electrical stimuli resulting from irreparable damage to motor neurons.

## 3. Conclusions

This study introduces an innovative approach focused on NaBC1 activation for the regeneration of muscles affected by ALS. Our findings demonstrate that the localized application of injectable borax-loaded alginate hydrogels in the quadriceps muscles of SOD1^G93A^ mice triggers NaBC1 activation, leading to enhanced muscle regeneration and repair processes. Following treatment, we observed a significant reduction of muscle inflammation, resulting in a tangible improvement in motor function and extended survival of mice. The efficacy of the treatment is further evidenced by a decline in muscle atrophy, as indicated by the decrease in slow type I fibers, diminished toxic SOD1 protein levels, and lower fibronectin levels. These outcomes, coupled with the substantial increase in active muscle satellite cells, the upregulation of essential muscle integrins, muscle trophic factors such IGF/IGFR and myogenic regulatory factors affirm the treatment’s effectiveness in promoting muscle growth, preventing muscle degeneration, and modulating inflammatory mechanisms crucial for muscle health via activation of the IGF–Akt–mTOR axis pathway. Notably, our study reveals that the activation of local muscle repair mechanisms induces retrograde neuroprotection by modulating neuro-inflammation, a key player in motor neuron death, ultimately preserving motor neurons.

In summary, this research provides compelling evidence supporting the involvement of muscle tissue in ALS pathology, emphasizing the potential of therapies targeting skeletal muscle for treating ALS. While ALS research typically focuses on motor neurons, the health of the muscles is paramount for overall motor function. Preserving muscle integrity through NaBC1 activation has the potential to generate neuroprotection. We propose a novel approach to decelerate neuromuscular degeneration in ALS, which may complement existing therapies targeting motor neuron inflammation.

## 4. Experimental section

### 4.1 Material substrates

#### Alginate-based hydrogel preparation

Ultrapure sodium alginate (Pronova^TM^ UP LVM, Novamatrix), was dissolved in 1% D-mannitol aqueous solution (Sigma-Aldrich) at a concentration of 1.5%. Borax (B) was dissolved in alginate solutions at 0.16 M concentration. Alginate solution was next filtered through a 0.22 µm pore Minisart Syringe Filter (Sartorius). For gelation, 2.7 mL of the alginate solution was mixed with 60 µL of 1.22 M CaSO_4_·2H_2_O (Sigma-Aldrich) through two Luer Lock syringe (BS Syringe) connected with a Fluid Dispensing Connector (Braun). Alginate and CaSO_4_·2H_2_O were mixed 10 times until complete homogenization. To retard the gelation time and produce injectable hydrogels, 60 µL of 0.5 M Na_2_HPO_4_·2H_2_O (Panreac) was added to the cross-linking reaction.

#### Borax-release determination

The *in vitro* release of borax previous to the animal experimentation was performed by immersing the alginate-based hydrogel (ALG and ALG-B), prepared with Na_2_HPO_4_ at 0.5 M, in 1 mL of DPBS with Ca^2+^ and Mg^2+^ (Gibco). Hydrogels were mixed as described above and performed with identical mass and diameter (6 mm) using agarose 2% (w/v) molds. Release studies were performed simulating an 8-day cell culture in a humidified atmosphere at 37 °C and 5 % CO_2_. Aliquots consisting of the total amount of immersing liquid (1 mL) were removed from the plates after diverse time points. The reaction of the borax present in the collected aliquots with azomethine (Sigma) in an acid medium (KAc/HAc buffer pH 5.2) originates a colorimetric reaction measured at 405 nm in a Victor III (Perkin Elmer) device. Standards for calibration were prepared at concentrations of 0, 0.1, 0.25, 0.5, 1, 1.5, 2.5, 5, 10, 25, 50, and 500 mg mL^−1^ of borax, using 40 µL aliquots from the original standard solutions for colorimetric reactions.

### 4.2 *In vivo* SOD1 mouse model

The experimental studies with the SOD1 mice model have been conducted according to Spanish and European regulations on the use and treatment of animals (RD 223/88, RD 53/2013, and 10/13/89 OM) and the ETS-123 European Convention on the protection/welfare of vertebrate mammals used in research. Specific authorization for the procedures was obtained (permit number 2020/VSC/PEA/0182), by the Animal Experiment Committee at Universitat Politècnica de València to ensure the avoidance of unnecessary distress, pain, harm, or suffering.

The experimental mice were housed under 12 h light / dark cycles at 23-25 °C with a relative humidity of 55 %. Food and water were available *ad libitum*. Animals were sacrificed by CO_2_ inhalation.

100 µL of injectable borax-loaded ALG-based hydrogels (ALG-B, 6 mg of borax/injection) were subcutaneously injected in both quadriceps muscles at 68, 82, 96, and 110 days old *B6SJL-Tg(SOD1-G93A)1Gur/J* ALS mice. Two different experiments were performed in parallel, differing in their endpoint: i) mice were sacrificed at 115 days old for muscle and nerve evaluation and ii) mice were sacrificed at their final endpoint determined as the mousés inability to flip over within 30 seconds in the supine position (ranging between 120-146 days) for evaluation of survival (SV).

For the first experiment, three different groups of male and female mice randomly distributed were evaluated: 6 animals injected with 100 µL of saline solution as an ALS control, 6 animals with empty ALG hydrogels, and 6 animals with ALG-B hydrogels. For the survival experiment (SV), two different groups of male and female mice randomly distributed were evaluated: 10 animals injected with 100 µL of saline solution as an ALS control and 10 animals with ALG-B hydrogels.

Changes in the body weight and motor impairment in the behavioral four-limb hanging test have been followed during the total duration of the experiments in all animals following Treat-NMD Neuromuscular Network Standard Operation Procedures (SOP, treat-nmd.org).

After euthanasia, different tissues were extracted:

- Both muscle quadriceps were removed and immediately fixed in cold isopentane (−150 ° to −160 °C) (PanReac Applichem) for 1 min, and subsequently immersed in liquid nitrogen for 2 min. Samples were stored at −80 °C until sectioning.
- The spinal cord was removed and immediately fixed in formaldehyde 4 % for 24 h at 4 °C, followed by cryopreservation in 30 % sucrose, inclusion in a freezing compound (OCT), and further stored at −80 °C until sectioning.

### 4.3 Histology and staining

Frozen muscle and spinal cord samples were sectioned using a cryostat at – 25/30 °C. 10-15 µm sections were placed on polarized slides (Superfrost, Thermofisher). Muscle histoarchitecture was evaluated by Hematoxylin-eosin staining. Muscle inflammation and motor neuron identification were determined by Toluidine Blue staining. All staining was performed following standard procedures. After sample dehydration with graded ethanol series and clearing with xylene, they were mounted with a xylene-based mounting medium (Entellan, Electron Microscopy Sciences). Images were captured with a bright-field microscope (Nikon Eclipse 80i) at different magnifications.

### 4.4 Immunofluorescence assays

Immunofluorescence assays were performed using cryopreserved slides containing sectioned samples that were air-dry for 15 minutes at room temperature. Samples were then permeabilized with DPBS/0.5 % Triton x-100 at room temperature for 5 min, next blocked in 2 % BSA/DPBS for 1 h at 37° C. Samples were then incubated with primary antibodies (Table 1**Error! Reference source not found.**) diluted in a blocking buffer overnight at 4 °C. The samples were then rinsed twice in DPBS/0.1 % Triton X-100 and incubated with the secondary antibody (Table 2) at room temperature for 1 h. Finally, samples were washed twice in DPBS/0.1 % Triton X-100 before mounting with Vectashield containing DAPI (Vector Laboratories) and images were captured under an epifluorescence microscope (Nikon Eclipse 80i) at different magnifications.

**Table 1.** Primary antibodies used for immunofluorescence assays.

| Target | Brand | Reference | Dilution |
| --- | --- | --- | --- |
| FN | Sigma-Aldrich | F0916 | 1:100 |
| GFAP | Abcam | ab7260 | 1:5,000 |
| IL-6 | ThermoFisher | P620 | 1:200 |
| PAX7 | Invitrogen | PA1-117 | 1:200 |
| Slow Skeletal Myosin Heavy Chain | Abcam | Ab234431 | 1:200 |
| SOD1 | Abcam | Ab51254 | 1:200 |

**Table 2.** Secondary antibodies used for immunofluorescence assays.

| Target | Brand | Reference | Dilution |
| --- | --- | --- | --- |
| Alexa Fluor 488 anti-mouse | Invitrogen | A-11029 | 1:700 |
| Alexa Fluor 555 anti-rabbit | Invitrogen | A-21428 | 1:700 |
| Alexa Fluor 555 anti-chicken | Invitrogen | A-21437 | 1:700 |

### 4.5 Gene expression analysis

Total RNA was extracted following the manufactureŕs instructions and under RNase-free conditions. Quadriceps muscle tissue (100 mg) from the different animal groups (ALS, ALG, ALG-B) was homogenized in 1 mL TRIZOL (Fisher Scientific) solution using a tissue Homogenizer 850 (Fisher) following 40 s cycles at 18.000 rpm. RNA quantity and integrity were measured with a NanoDrop 1000 (ThermoScientific). Then 2-5 µg of RNA were reverse transcribed using the Superscript III reverse transcriptase (Invitrogen) and oligo dT primer (Invitrogen). Real-time qPCR was performed using Sybr select master mix and 7500 Real-Time PCR system from Applied Biosystems. The reactions were run in triplicate for technical replicas while biological replicas were 6 mice in the case of pre-symptomatic animals and 10 mice in the case of symptomatic animals. The primers used for amplification were designed from sequences found in the GenBank database and are indicated in Table 3.

**Table 3.**
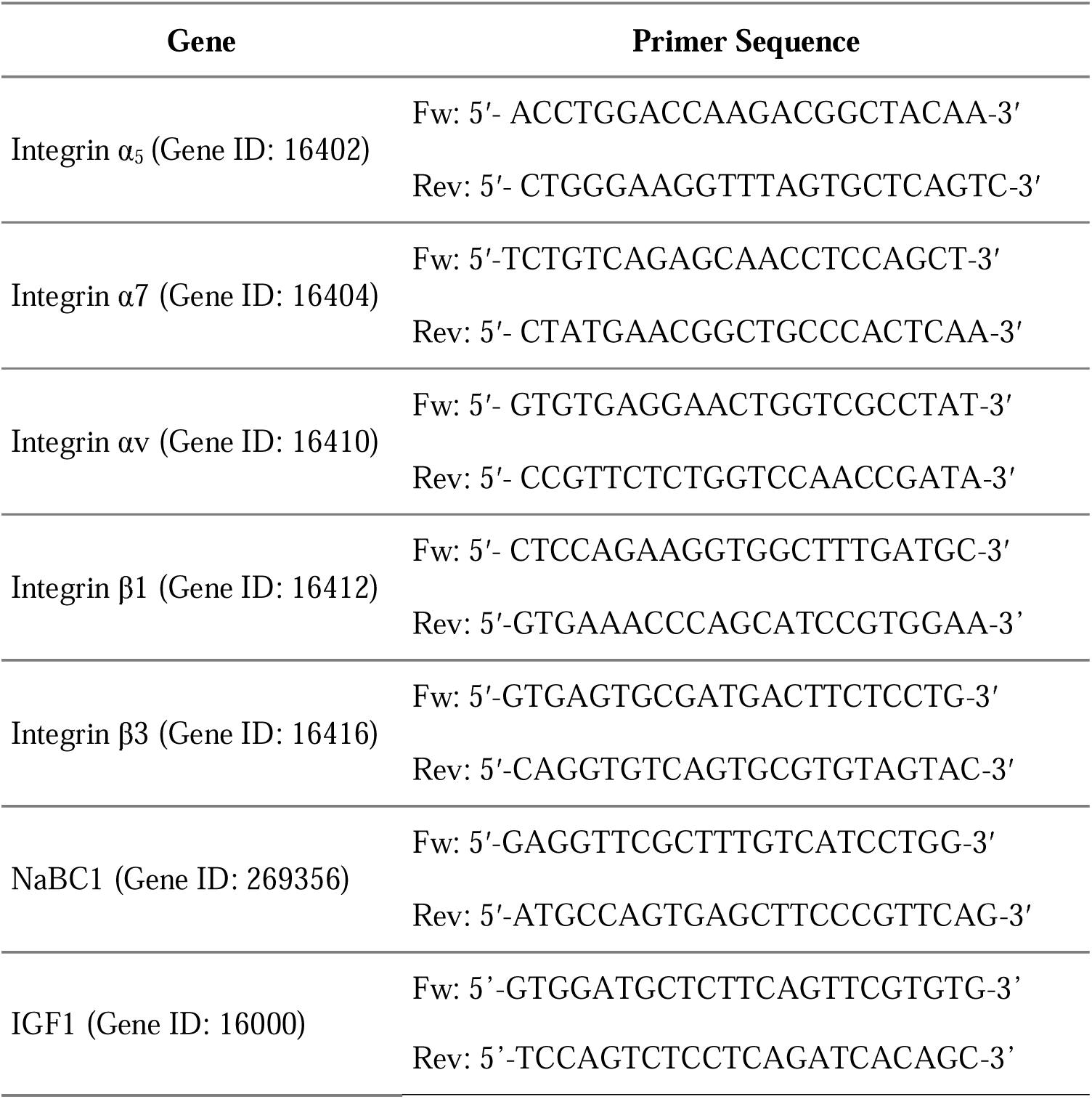
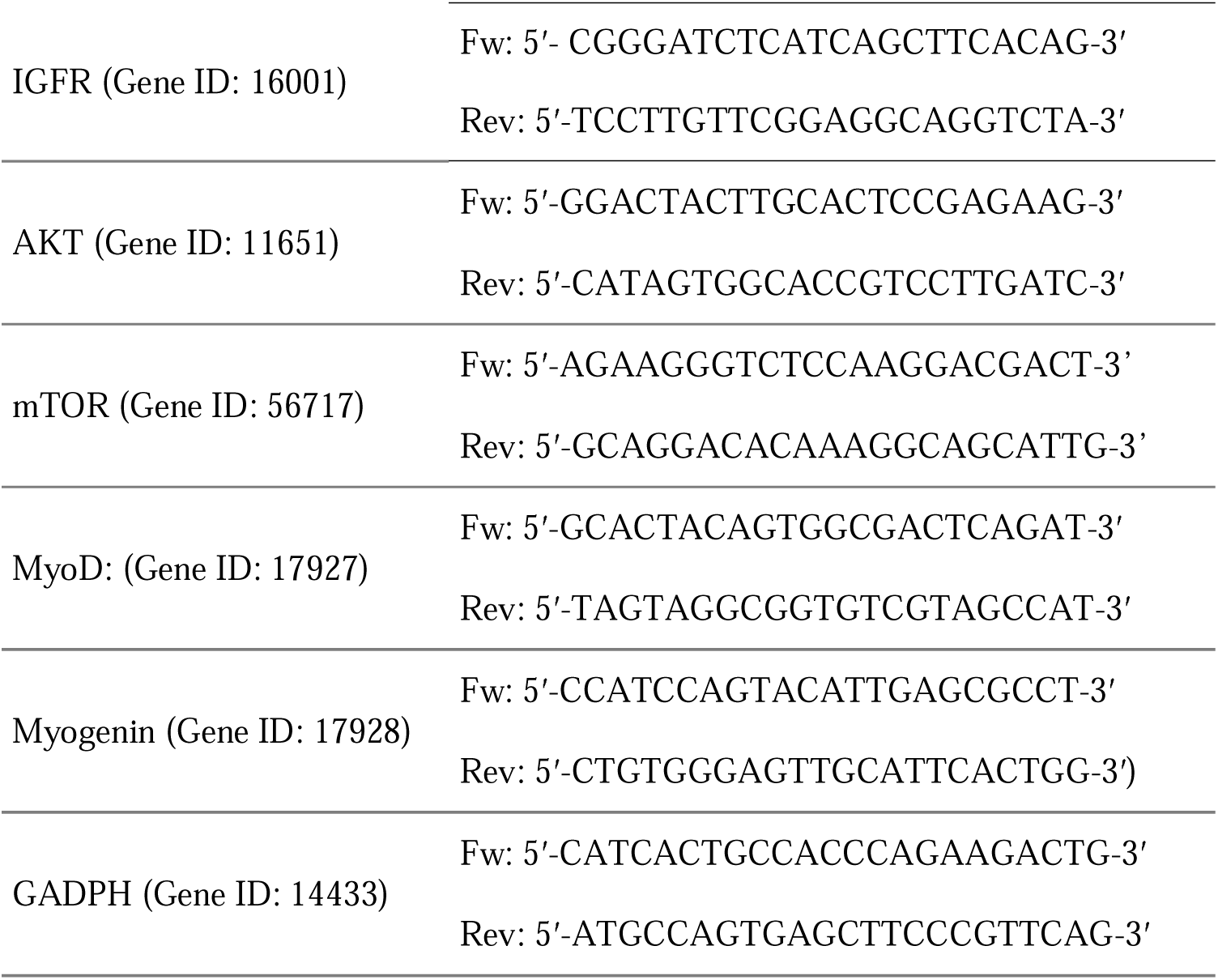
Primer sequences are used for gene expression analysis.

### 4.5 Image analysis

Muscle sequential images obtained after staining were stitched (Image Composite Editor) and segmented using Cellpose 2.0 a generalist algorithm for segmentation, allowing the identification of cell boundaries and the analysis of all fibers of a muscle cross-section (2,000-5,000 fibers). This method for analyzing the entire muscle guarantees unbiased results. Muscle histological features were identified manually and quantified using image analysis software ImageJ following Treat-NMD Neuromuscular Network Standard Operation Procedures (SOP, treat-nmd.org). The staining intensity of immunofluorescence images was quantified by ImageJ software.

### 4.6 Statistics and Reproducibility

For statistical analysis GraphPad Prism 8.0.2 has been used. Data were reported as mean ± standard deviation. Normal distribution of the data was established using the D’Agostino Pearson omnibus test. Results were analyzed by one-way ANOVA test with Tukeýs multiple comparison test (p = 0.05) when 3 or more groups were compared. Pairs of samples were compared using unpaired t-tests, applying Welch’s correction when necessary. A 95% confidence level was considered significant and is indicated by (*) p < 0.05, (**) p < 0.01, (***) p < 0.001, and (****) p < 0.0001.

## Acknowledgments

Funding: PR acknowledges support by grant PID2021-126012OB-I00 funded by MCIN/AEI/10.13039/501100011033 and by ERDF a way of making Europe, and by CIBER (CB06/01/1026). JGV acknowledges the funding from the European Union-NextGenerationEU program. PR acknowledges gratefully the charitable funding provided by the ALS patient LEP and her family and the help provided by CGR, JVH, and PMN.

