## supplementary information for "Injectable borax-loaded alginate hydrogels reduce muscle atrophy, modulate inflammation, and generate neuroprotection in the SOD1^G93A^ mouse model of ALS via activation of the IGF–Akt–mTOR axis pathway"

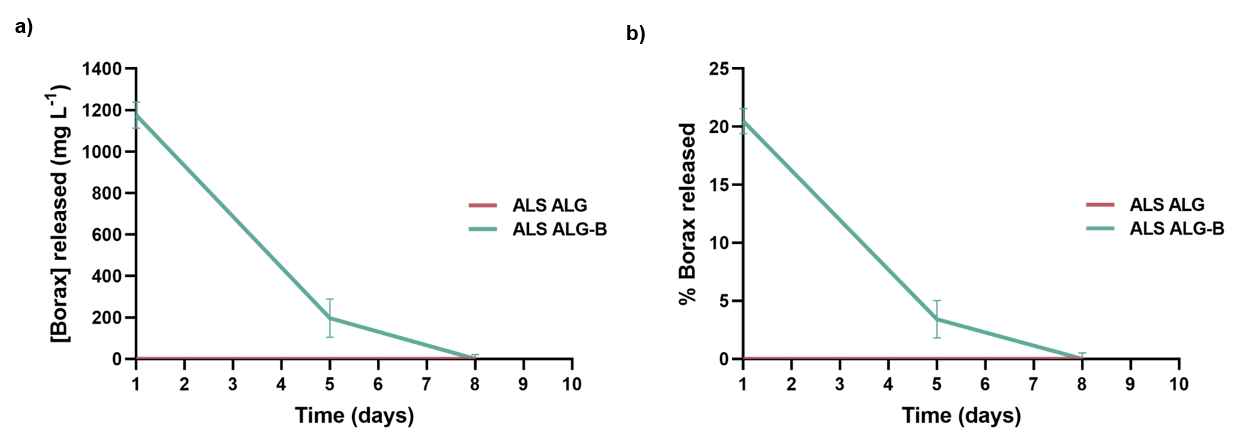
**Supplementary information**

**Figure S1. Evaluation of borax-release.**

a) Long-time non-cumulative borax-release assay of 6 mm injectable ALG-B hydrogels after, 1, 5, and 8 days, expressed in mg L^-1^.

b) Percentage of borax released from alginate hydrogels, indicating that approximately 80 % of borax remains retained within the hydrogel.

Statistics are shown as mean ± standard deviation.

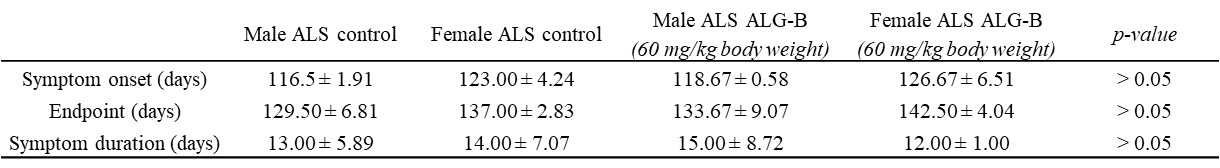
**Table S1**. **Analysis of symptoms during the disease progression.**

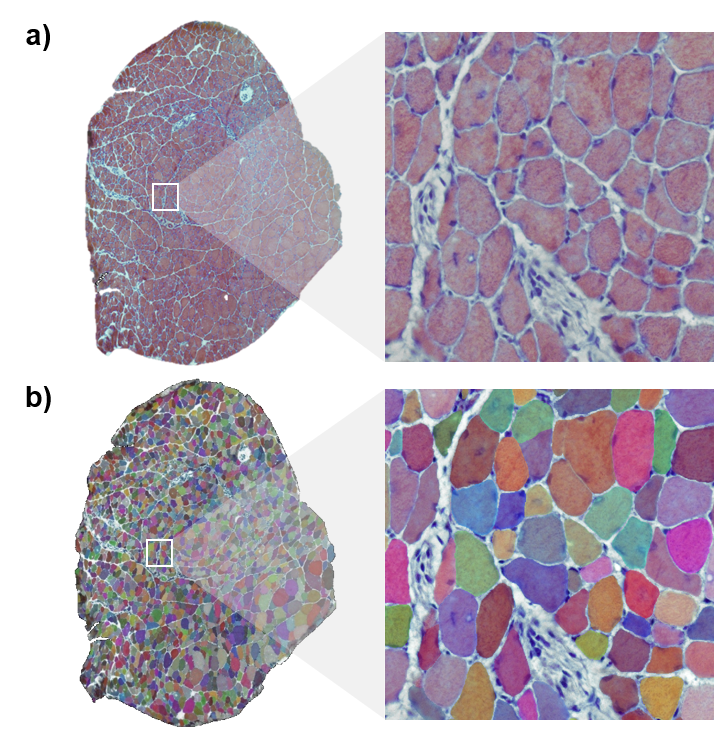

**Figure S2. Segmentation of SOD1^G93A^ complete muscle sections after Hematoxylin-eosin staining for individual fiber quantification.**

a) Total quadriceps histological section from SOD1^G93A^ mice stained by hematoxylin-eosin.

b) Obtained mask of total quadriceps histological section after segmentation with CellPose software. This method allows quantification of the total amount of fibers present in the complete muscle section.

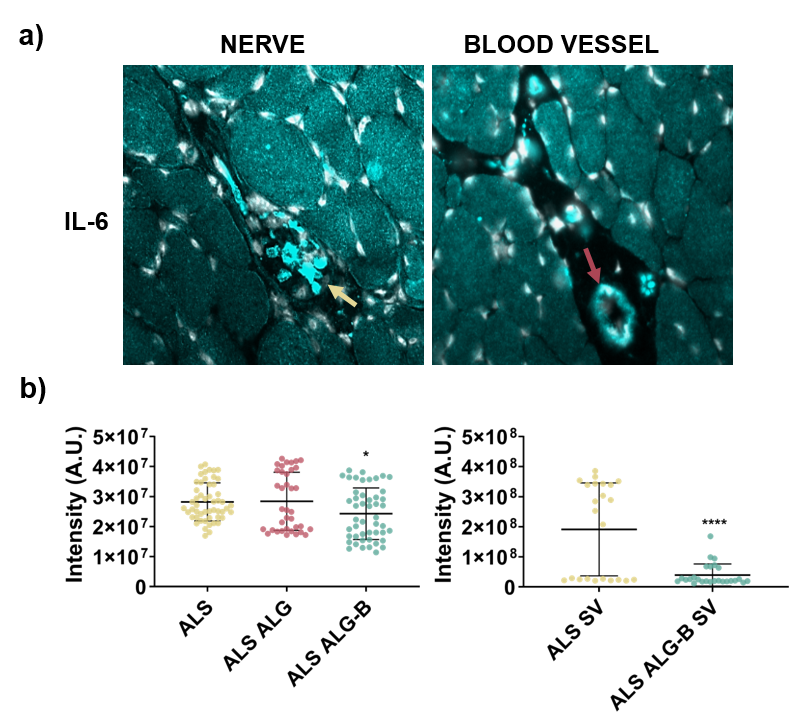

**Figure S3. Evaluation of the inflammatory cytokine IL-6 in symptomatic and terminal-stage SOD1G93A mice.**

**a)** Immunofluorescence detection of IL-6 (Cyan) in nerve and blood vessels. Nerve and blood vessel structures are indicated by yellow and red arrows respectively.

**b)** Quantification of IL-6 staining intensity in symptomatic and terminal-stage mice. Treatment with ALG-B in both groups decreases inflammation.

Statistics are shown as mean ± standard deviation. For comparison between the three groups (ALS, ALG, ALG-B), data were analyzed by an ordinary one-way ANOVA test and corrected for multiple comparisons using Tukey´s correction analysis (p = 0.05). For comparisons between the two groups (ALS SV, ALG-B SV), data were analyzed by an unpaired t-test applying Welch´s corrections (p = 0.05). ****p < 0.0001, *p < 0.05.
